# Connexin 43 mediated collective cell migration is independent of Golgi orientation

**DOI:** 10.1101/2023.05.03.539214

**Authors:** Madhav Sharma, Suvam Mukherjee, Archana Kumari Shaw, Anushka Mondal, Amrutamaya Behera, Jibitesh Das, Abhishek Bose, Bidisha Sinha, Jayasri Das Sarma

**Author notes:** equal contribution; shared first author (Madhav Sharma, Suvam Mukherjee, Archana Kumari Shaw).

## Abstract

Cell migration is vital for multiple physiological functions and is involved in the metastatic dissemination of tumour cells in various cancers. For effective directional migration, cells often reorient their secretory traffic towards the leading edge by reorienting the Golgi apparatus. However, conflicting results on the positioning of the Golgi relative to the migrating cell raise questions about its regulation. Herein, we address the role of gap junction protein Connexin 43, which connects cells allowing the direct exchange of molecules and also controls the distribution of microtubules. We utilized HeLa WT (wild-type cells lacking Connexin 43) and HeLa 43 cells (stably expressing Connexin 43). We found that functional Connexin 43 channels affected Golgi morphology and reduced the ability of reorientation of Golgi (towards the leading edge) during cell migration. Although Connexin 43 also reduced migration velocity, the front displayed enhanced coherence in movement, implying an augmented collective nature of migration compared to HeLa WT cells lacking the cell-cell connectivity evidenced in HeLa 43. The increase in vimentin and basal actin in HeLa 43 shows that Connexin 43 expression alters the cytoskeleton. Non-invasive measurement of basal membrane height fluctuations revealed that HeLa 43 had a lower membrane tension and a more uniform intra- and inter-cellular distribution than HeLa WT, similar to their coherence in migration. We, therefore, propose that the reduced Golgi reorientation in HeLa 43 might be linked to lesser dependence on directed secretory trafficking, which is believed to be required to buffer the increasing tension caused by migration.

## Introduction

Cell migration is a fundamental process that has a pivotal role in early life when embryogenesis occurs^1^. It is also essential for many physiological functions, e.g., immune surveillance, angiogenesis, and wound healing^2^. Moreover, cell migration plays an essential role in pathophysiological processes such as tumour growth and metastasis and vascular remodeling in various diseases^3^. Directional migration is achieved by establishing front–back polarity in migrating cells^4^. In addition to the asymmetric organization of the cytoskeleton, which is achieved through recruitment and the local synthesis of proteins, polarized cells reorient their secretory traffic towards the direction of migration^5,6^. Such exocytic cargos, which originate from the Golgi, contain additional membrane, cell-surface receptors, and extracellular matrix components that are required to maintain the leading edge^7,8^. Several proteins are involved in controlling cell migration or its modulation^9^. Among them, connexins, which are classically considered to act as gap junction channel-forming proteins, are known to play an essential role in determining directional migration^10^. Gap junctions are specialized cell junctions that contain hydrophilic membrane channels that allow the passive diffusion of ions and small molecules between cells^11^.

All connexins exhibit a conserved protein structure comprising four transmembrane domains and a cytoplasmic tail localized carboxy terminus with critical regulatory functions^12,13^. Connexin 43 (Cx43) is an essential member of the connexin family, ubiquitously expressed across several tissues in a cell-specific manner^14,15^. Its function, however, is not limited to the transport of small molecules. Cx43 plays a significant role in cell migration and polarity by regulating the microtubule network^10^. Cx43 is synthesized in the ER as an unglycosylated four membrane-spanning monomeric transmembrane protein. Connexin 43 subunits are post-translationally assembled as hexamers at the trans-Golgi network as Cx43 hexamers are transported in vesicles along the microtubular conduit to reach the cell surface, where they form hemichannels (connexon) and subsequently associate with another connexon to form GJ channels^14^. Studies have shown that polarized Golgi is vital for normal developmental processes and directed secretion of extracellular factors, responsible for effective cell migration. In a polarized cell, Golgi is pericentrosomally oriented towards the direction of migration^16^. Intracellular remodeling of Golgi happens through the microtubule, with the latter tethering Golgi at the polarization region^17^.

Leader cells are the first to explore the tissue microenvironment during cell migration^18^. Thus, they contribute to the navigation of follower cells. For the leader cells, directing the Golgi towards any direction of the mechanical void makes the Golgi directed towards the scratch in a scratch assay^19^. By secretion of extracellular matrix (ECM), leader cells create a substratum for the followers and make it easy for them to migrate^20^. Follower cells also influence the leader cells through the cell surface molecules like cadherins^21–23^, ephrins^24–26^, members of the polar cell planarity complex, i.e., Frizzled^27^, WNT^28^ and, PTK7^29^ and syndecan 4^30^, thereby encouraging directed migration of leader cells. Thus, in collective cell migration, leaders and followers influence each other through their continuous, directed motion^31,32^. In systems like keratocyte cell sheets, a lateral connection between neighboring cells and actin cables across multi-cellular length scales coordinate the coherent movement of the front^33^.

In contrast, in single cells, the coherence of the lamellipodia extension is responsible for converting random migration to directed cell migration^34,35^. A front-back gradient of membrane tension has been reported and is believed to be critical to channel forces in one direction^36^. Directing secretory machinery (of the Golgi apparatus) towards the leading edge assures timely membrane delivery through exocytosis and maintenance of the front tension, which would otherwise keep increasing due to actin polymerization forces^37^. Despite this role of the Golgi apparatus, multiple studies report Golgi to be not necessarily oriented towards the leading edge of migrating cells^38^. There exists a clear gap in our understanding of the dependency of migration on Golgi re-orientation. In particular, enhancing cell-cell communication through GAP junctions can fundamentally alter information flow between cells, reducing mechanical signals’ impact, and increasing chemical connectivity. It remains unexplored if cells depend on front-back tension gradient and Golgi re-orientation when cell-cell communication is established.

Thus, although Golgi orientation in migrating cells plays a crucial role, the underlying mechanism and its correlation with the presence of functional GAP junctions need to be adequately evaluated. Towards this, our study using Hela WT (Cx43-/-) and Hela 43 (Cx43+/+) demonstrated that expression of connexin 43 is associated with morphological alteration of the Golgi apparatus in HeLa WT cells. Interestingly, we also observed the expression of vimentin, mesenchymal cell protein, and DJ-1 protein, a known antioxidant protein indicating more inclination towards the mesenchymal character, suggesting that in the tussle between the presence of gap-junctions and mesenchymal characteristics, functional gap junctions in HeLa WT cells slowed down the collective cell migration. The basal actin was imaged using total internal reflection fluorescence (TIRF) microscopy and revealed how Connexin-43 affects the migrating cell surface. Interference reflection microscopy (IRM) was used to assess the concomitant changes in the mechanics of the basal membrane in the presence of Connexin 43 that might be linked to the slowed and coherent migration.

## Material and Methods

### Cells and Cell lines

HeLa WT cells obtained from ATCC and maintained in Dulbecco’s Modified Eagle Medium supplemented with 10% Fetal Bovine serum (Invitrogen) and 1% penicillin-streptomycin (Gibco). HeLa WT cell line stably expressing Connexin 43, referred to as HeLa 43, were generated as previously described (JDS,2002)^39^. HeLa 43 cells were maintained in the same medium mentioned above with additional supplementation of G418 (Sigma, A1720) at a final concentration of 500μg/ml.

### Lucifer yellow dye transfer

The gap junction functional coupling between HeLa WT and HeLa 43 cells was determined using scrape loading of Lucifer yellow (LY) in confluent monolayers of HeLa WT and HeLa 43 as described previously^40^, with minor modifications. HeLa WT and HeLa 43 were scrapes loaded with PBS containing 4 mg/ml Lucifer yellow CH (Sigma, Saint Louis, MO). After a 1-min incubation, the LY solution was removed from the culture, the culture was washed thoroughly with PBS, and respective fresh culture medium was added. The distance of LY spread from the scrape-loading point to neighboring cells was imaged using an inverted microscope with a Hamamatsu Orca-1 charge-coupled device (CCD) camera, and the distance spread was measured using ImageJ software.

### Cell migration assay

Scratches were created on a confluent monolayer of HeLa WT and HeLa 43 with thorough washing with PBS to remove non-adherent cells, after which cell-specific media was added. Cells were monitored for migration capacity at 0, 4, 8 and 12 hours using a bright field microscope. The culture medium was supplemented with 0.2 μM of Hoechst solution for immunofluorescence studies. Results were analyzed using ImageJ software and interpreted as described in supplementary materials.

### Immunofluorescences

Immunofluorescence studies were done according to a protocol described previously^41^, with minor modifications. For standard immunofluorescence, Cells were plated on etched glass coverslips and fixed with 4% paraformaldehyde (PFA). Permeabilization was done with phosphate-buffered saline (PBS) containing 0.5% Triton X-100 and blocked with blocking solution (PBS containing 0.5% Triton X-100 and 2.5% heat-inactivated goat serum (PBS-GS)). The cells were incubated with primary antisera diluted in a blocking solution for 1 h, washed, and then labeled with secondary antisera diluted in a blocking solution. Then cells were visualized using a Zeiss confocal microscope (LSM710). Images were acquired and processed with Zen2010 software (Carl Zeiss).

### Membrane protein isolation and analysis

Monolayer of HeLa WT and HeLa 43 cells were washed, then harvested in PBS containing protease inhibitor cocktail (complete protease inhibitor; Roche, Mannheim, Germany) and phosphatase inhibitors (1 mM NaVO4 and 10 mM NaF), and then passed through a Dounce homogenizer 100 times^42^. The homogenate was centrifuged at 500g for 5 min using an Eppendorf 5415 R centrifuge. The resulting supernatant was centrifuged at 100,000g for 30 min using a Beckman Optima Max ultracentrifuge to obtain a membrane-enriched pellet. To analyze total membrane connexin expression, this pellet was resuspended in RIPA buffer and subjected to immunoblot analysis for Cx43.

For detergent solubilization studies, the membrane-enriched pellet was resuspended in PBS with protease and phosphatase inhibitors at 4°C containing 1% Triton X-100 and then incubated for 30 min at 4°C. Care was taken to ensure the samples were not warmed to RT during extraction. The sample was centrifuged at 100,000 g for 30 min and separated into Triton X-100 soluble supernatant and insoluble pellet fractions^43^. The soluble fraction was then diluted into RIPA buffer, while the insoluble fraction was initially resuspended in PBS with 1% Triton X-100 before dilution. Equal volumes of soluble and insoluble fractions were probed for Cx43 by Immunoblotting.

### Immunoblot Analysis

For immunoblotting, samples were resolved by SDS-PAGE using a 12% polyacrylamide gel, transferred to polyvinylidene difluoride (PVDF) membranes using transfer buffer (25 mM Tris, 192 mM glycine, and 20% methanol), and blocked for 1 h at RT using blocking solution (5%, wt/vol, powdered milk dissolved in TBST or Tris-buffered saline containing 0.1%, vol/vol, Tween 20). The samples were then incubated overnight at 4°C in primary antisera diluted in blocking solution, followed by washes with TBST and then a 1-h incubation of horseradish peroxidase (HRP)-conjugated secondary IgG in blocking solution. The immunoblots were washed in TBST, and then immunoreactive bands were visualized using Super Signal WestPico chemiluminescent substrate (Thermo Scientific, Rockford, IL). Densitometric analysis of nonsaturated films was performed using Bio-Rad Quantity One analysis software (Hercules, CA) or using a Syngene G: Box ChemiDoc system and GENESys software

### Interference Reflection microscopy

For obtaining membrane fluctuation and fluctuation tension, interference reflection microscopy (IRM) was performed in an onstage 37° C incubator (Tokai Hit, Japan). An inverted microscope (Nikon, Japan) equipped with a 100 W mercury arc lamp, an interference filter (546 ± 12 nm), a 50-50 beam splitter, a 60X water-immersion objective (1.22 NA), and an sCMOS camera (Hamamatsu, Japan) was used. Imaging was performed with 50 ms exposure time capturing a total of 2048 frames for any single measurement.

### Total Internal Reflection Fluorescence Microscopy

For assessing basal actin in migrating cells using TIRF, cells were grown into a monolayer and were scratched to create a wound-like region. Cells were subsequently incubated for 3 hours and 45 mins, followed by applying CellMask Orange Actin tracking stain (Invitrogen, US) at a 0.5 μM. Cells were imaged after 15 mins of incubation in the live actin stain.

Imaging was performed using an inverted microscope (Olympus, Japan) with a 100x 1.49 NA oil immersion TIRF objective using a 561 nm laser. The exposure time of 200 ms and a final pixel size of 65 nm was maintained throughout.

### Quantification and Statistical Analysis

Calibration and analysis of IRM were performed as described previously^44^. Briefly, beads (60 μm diameter polystyrene beads; Bangs laboratories) were imaged on each day of experiments to find the intensity to conversion factor after comparison with IRM images of cells. Subsequently, pixel, where this conversion held, were identified and termed first branch regions (FBRs). Groups of 12×12 pixels (0.89×0.89 μm^2^) were used to sample single cells locally to recreate migrating cells’ tension landscape. Measurements of fluctuations only from FBRs were used for quantitative analysis. The standard deviation (SD) of relative heights (of 2048 frames) calculated at every pixel of an FBR (144 pixels) was averaged to find the SD_time_. To find the tension (fluctuation tension), the power spectral density of the fluctuations was obtained using MATLAB’s covariance method (autoregressive PSD estimation) and fitted with the Helfrich-based theoretical model also used earlier.

## Results

### Golgi orientation is random in the absence of migration

The Golgi apparatus can be distributed in different forms in cells. However, HeLa WT cells usually display a clustered distribution, as visible in the images of HeLa WT cells expressing mCherry-Golgi7 (**Fig. 1A**). To quantify its orientation (**Fig. 1B**), the Centre of mass (CM) of the image of the nucleus is joined to the CM of the Golgi to get the angle. Without any “front,” no particular direction can be used as a reference. Hence, the x-axis of the image is taken as the reference. The angle is measured for it (**Fig. 1B**). Once such angles are obtained for all cells in a particular dish (sharing a common x-axis), distributions do not show any angle favored. Pooling over multiple dishes from multiple trials (**Fig.1C**), the distribution was statistically not significantly different from a uniformly generated random numbers distribution.

**Figure 1.**
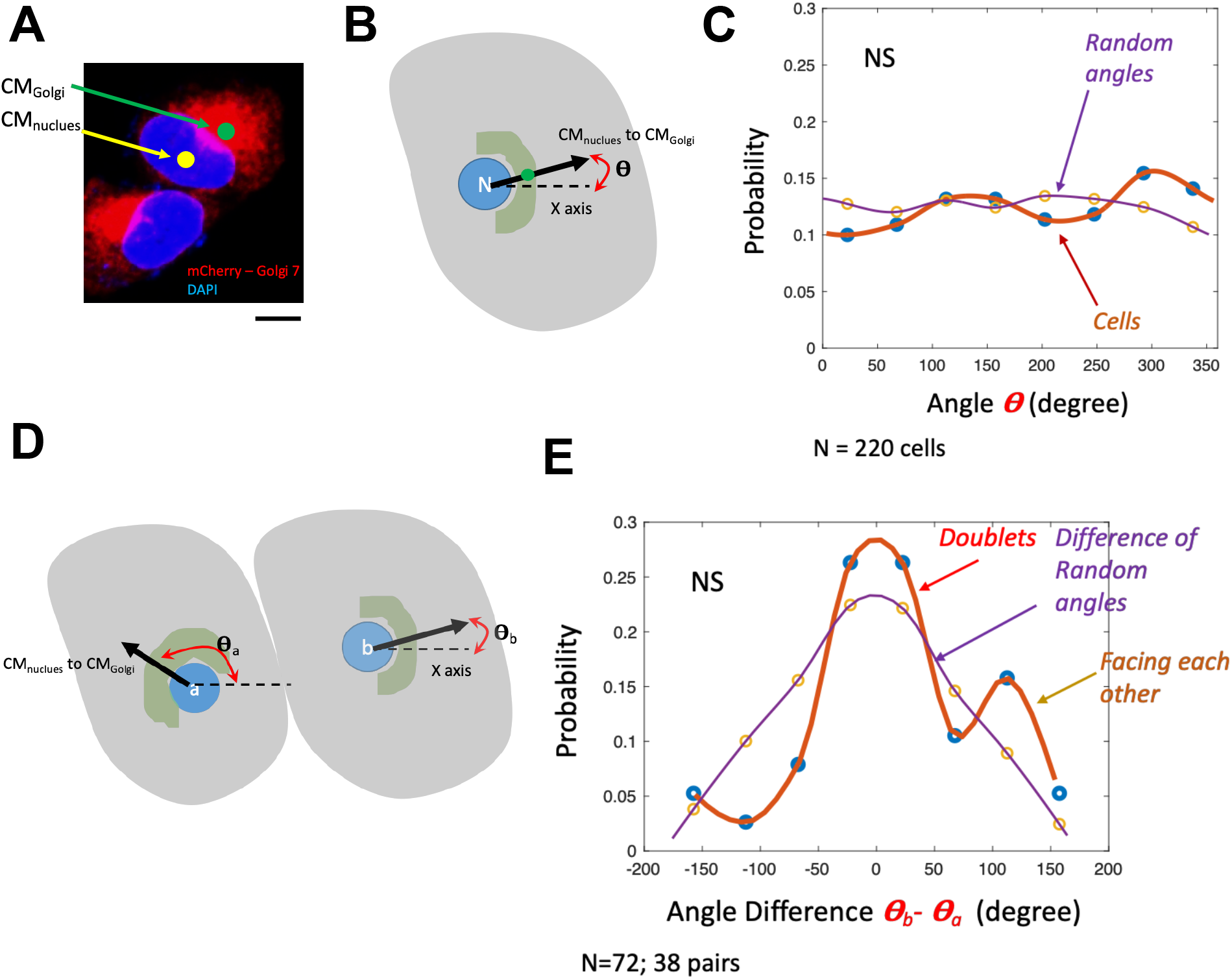
Quantifying Golgi’s orientation. (A) Representative image of cells expressing mCherry-Golgi 7 (red) and stained with DAPI (blue). Scale bar = 10 μm. (B) Schematic of angle measurement from CMs measured for nucleus and Golgi using image analysis (C) Angle distribution and its comparison to normally distrusted random numbers in the same range. No significant difference was found as expected. Wilcoxon rank-sum test was performed to test for significance. N=220 cells. D) Schematic representation of angle measurements for doublets (cell represented in gray, Golgi represented in green and nucleus represented in blue). For each cell, the angle (***θ***) subtended at the x-axis, by the line joining its CM_nucleus_ to CM_Golgi_ is first measured. Cell whose CM_nucleus_ is at a bigger x value (or cell is to the right) is termed as the “b”. (E) Distribution of ***θ***_***b***_ − ***θ***_***a***_ and distribution of difference of uniformly distributed random angles in the same range as observed for cells. Angle differences obtained from cells was not significantly different from difference of random angles as tested by Wilcoxon rank-sum test.

To understand if cell-cell interactions were enough to reorient Golgi – even in the absence of cues from the wound, we next analyzed images of cell “doublets” (**Fig. 1D**). The centroid of the binary image of the nucleus (marked as CM_nucleus_) was joined to the Centre of mass (CM) of the Golgi (calculated from the grayscale image taking account of intensity variations) to get the angle with respect to the x-axis of the image – same for all cells in a dish. For each cell, the angle (θ) subtended at the x-axis, by the line joining its CM_nucleus_ to CM_Golgi_ was first measured. The cell whose CM_nucleus_ was at a more considerable x value (or cell is to the right) was called the “b.” The distribution of θ_b_ − θ_a_ and the distribution of the difference between two uniformly distributed random angles in the same range was compared and statistically tested. Angle differences obtained from cells were not significantly different from those of random angles (**Fig. 1E**) as tested by the Wilcoxon ranksum test in MATLAB.

Thus, for doublets, although we find that when there is a slightly higher probability of finding cells facing each other than if they are random – however, the data remained statistically non-significant from simulated data mimicking the random direction of the cells. This established that measured orientations were not substantially influenced by how neighboring cells are arranged without any spatial cue, like a wound.

### Stacked Golgi apparatus in HeLa sensitive to Connexin 43

Golgi apparatus is the protein glycosylation and transport machinery of the cells and plays an important role in membrane trafficking. HeLa WT cells, which are epithelial carcinoma cells, were used to understand its role in cell migration. Golgi apparatus of Hela WT cells were characterized using Golgin 97 protein, predominantly present in Trans Golgi Network (TGN) and plays critical roles as a tethering molecule associated with tubulovesicular carriers during vesicle trafficking and in maintaining the Golgi integrity^45^. Immunofluorescence with anti-Golgin 97 antibodies indicates constricted Golgi in HeLa WT (**Fig. 2A, B**). Endoplasmic Reticulum of HeLa WT is also characterized using anti-Calnexin Antibody, a chaperone exclusively present ER, indicated endoplasmic reticulum is more conical shaped (**Fig. 2C**). We also showed that HeLa WT was negative for connexin 43 expression. To understand the role of connexin 43 in cell migration, Hela 43, HeLa WT cells stably expressing connexin 43 were used. Immunofluorescence data suggests that cell surface expression of connexin 43 is associated with diffused staining of Golgin 97 denoting altered Golgi structure. The Golgi appeared more crescent-shaped in HeLa 43 as opposed to constricted shaped in Hela WT, suggesting that exogenous expression of connexin 43 at the cell surface as well as intracellular retention, causes the altered distribution of Golgi stacks (**Fig. 2B**). Also, HeLa 43 cells displayed gap junction of connexin 43 in the form of puncta as shown (**Fig. 2B**). In addition to addressing the role of Connexin 43 in Golgi’s reorientation and migration, we also used Brefeldin A (BFA) to create ER stress and thereby convert a stacked Golgi to the distributed vesicular organization in the cell (**Fig. 2D**).

**Figure 2.**
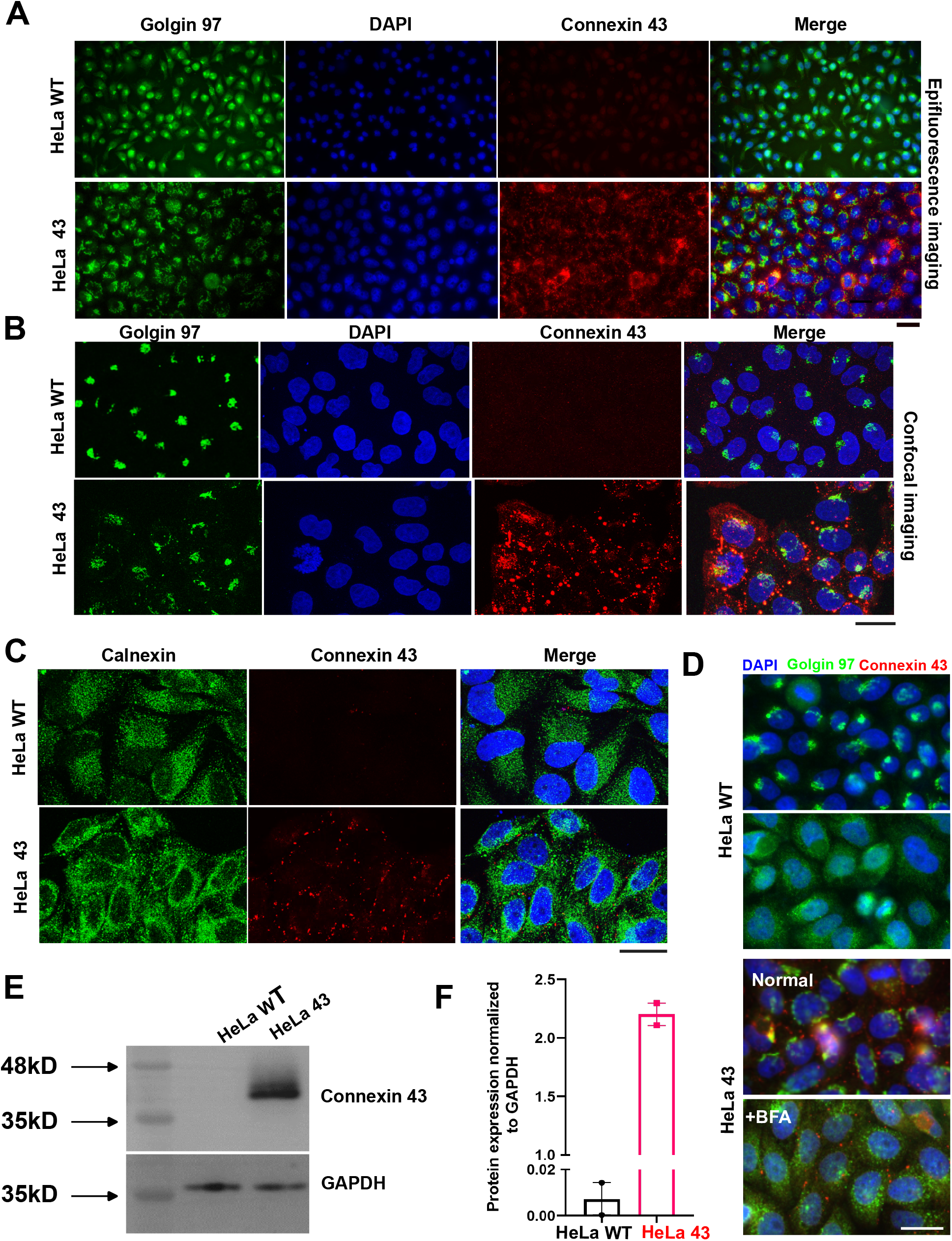
Characterizing HeLa WT and HeLa 43. (A) Epi-fluorescence images of HeLa WT and HeLa 43 highlighting Golgi and Connexin 43 distribution (B) Confocal images resolving better the altered morphology of Golgi in HeLa 43 (C) IF with Calnexin antibody reveals no particular difference in ER in the two cell lines. (D) BFA alters Golgi distribution (E) Westerns blots showing higher abundance of Connexin 43 in HeLa 43. (F) Quantification of western blot. A: Scale Bar = 100 μm; B-D: Scale Bar = 30 μm.

Immunoblot assay (**Fig. 2E, F)** using anti-Connexin 43 antibody confirms the presence of connexin 43. In contrast, HeLa WT was found to be negative for connexin 43 expression both in immunofluorescence and immunoblot assays confirmed by quantification (**Fig. 2F**).

### HeLa 43 forms functional gap junctions

To check the functional gap junction, the scrape loading/dye transfer technique was used, a simple, functional assay for the simultaneous assessment of gap junctional intercellular communication (GJIC) between adjacent cells in a large population of cells. As Cx43 channels are permeable to LY (<1 KD), a small molecular dye that moves from cells to the neighboring cells via gap junctions, the green color dye transfer in the HeLa 43 cells indicates the presence of functional gap junctions formed by connexin 43 as opposed to HeLa WT which does not establish functional gap junctions (**Fig. 3A**). Furthermore, Cx43 assembled into gap junction plaques is resistant to 1% Triton X-100 solubilization at 4°C, while remaining Cx43 dissolves in the Triton X-100 soluble fraction. Therefore, Triton X-100 solubilization was performed to determine the retention of Cx43 in the intracellular compartment instead of it being trafficked to the cell surface to form gap junction plaques. The membrane fraction was isolated from cells and was solubilized using 1% Triton X-100 at 4°C, separated into detergent-soluble and - insoluble fractions, and probed for Cx43. HeLa 43 showed that most Cx43 was pooled into the Triton X-100 insoluble fraction rather than the soluble fraction. Hence, the insoluble/soluble fraction ratio of Cx43 expression was significantly higher in HeLa 43 (**Fig. 3B**). This experiment demonstrated that a small amount of Cx43 retains in the intracellular compartment (triton soluble fraction). In contrast, a large fraction of Cx43 was observed in the insoluble fraction, which denotes that Cx43 forms insoluble gap junction plaque at the cell surface.

**Figure 3.**
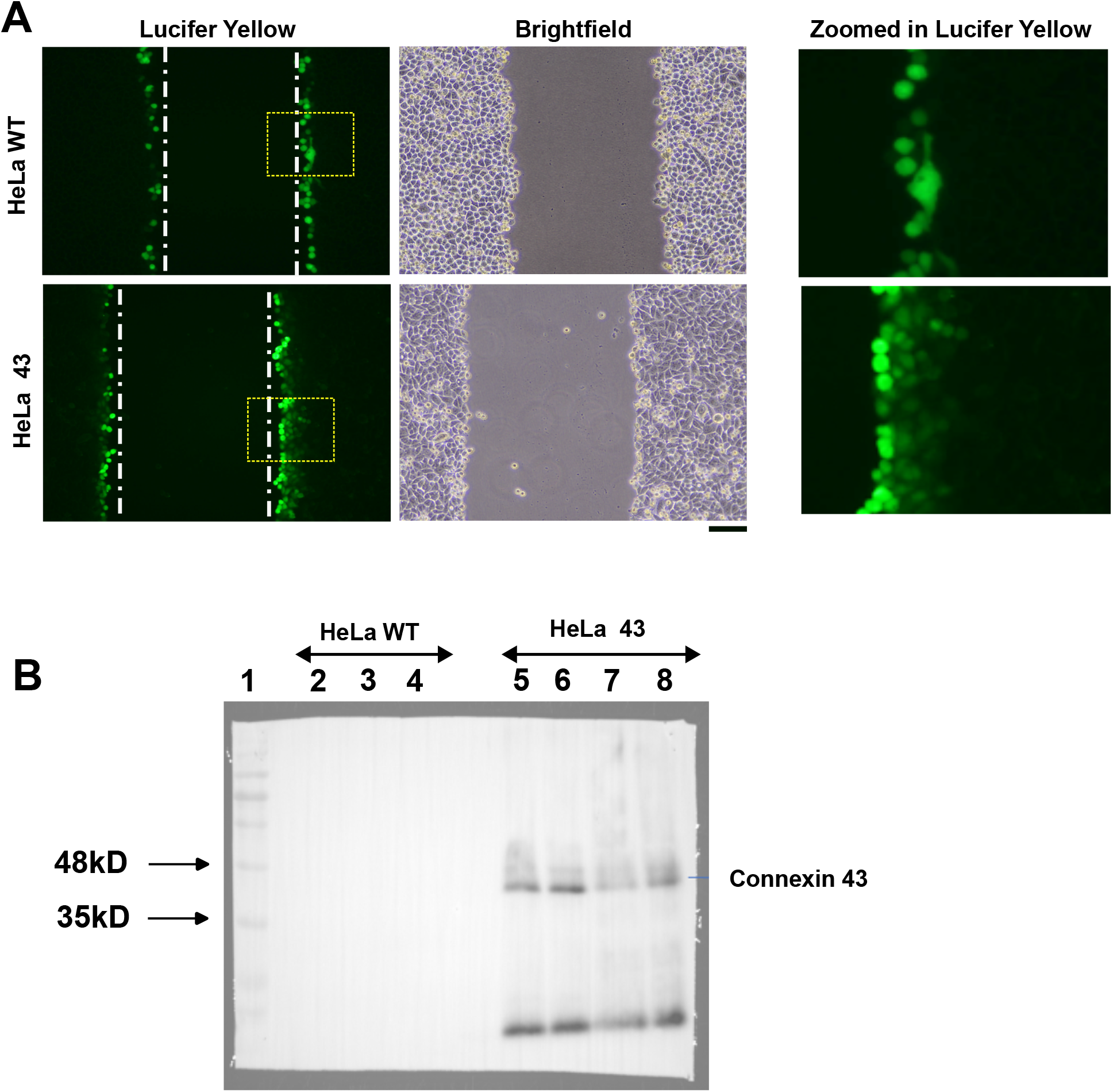
Evaluating gap junction functionality. (A) Scrape loading assay depicted with representative scrapes with LY shown in green. Vertical whit elines denote the scrape line. Rectangular ROIs outlinr regions expanded in the rightmost column (B) Comparison of Tx-solubilisation between HeLa WT and HeLa 43. 2 & 3, HeLa WT insoluble fraction; 4 HeLa WT soluble fraction; 5 & 6, HeLa 43 insoluble fraction; and 7 & 8, HeLa 43 soluble fraction. Scale bar = 100 μm.

### Golgi reorientation during migration is lower in HeLa 43

We next set up the scratch assay in which an asymmetry of neighbours would be introduced. To quantify the reorientation of Golgi as the migration progresses. Using immunofluorescence on samples fixed at different time points (**Fig. 4A**), we imaged the Golgi’s distribution in cells that face the wound (termed as ‘first line’ cells) as well as cells not directly exposed to the scratch but separated by a line of cells (termed ‘second line’ of cells).

**Figure 4.**
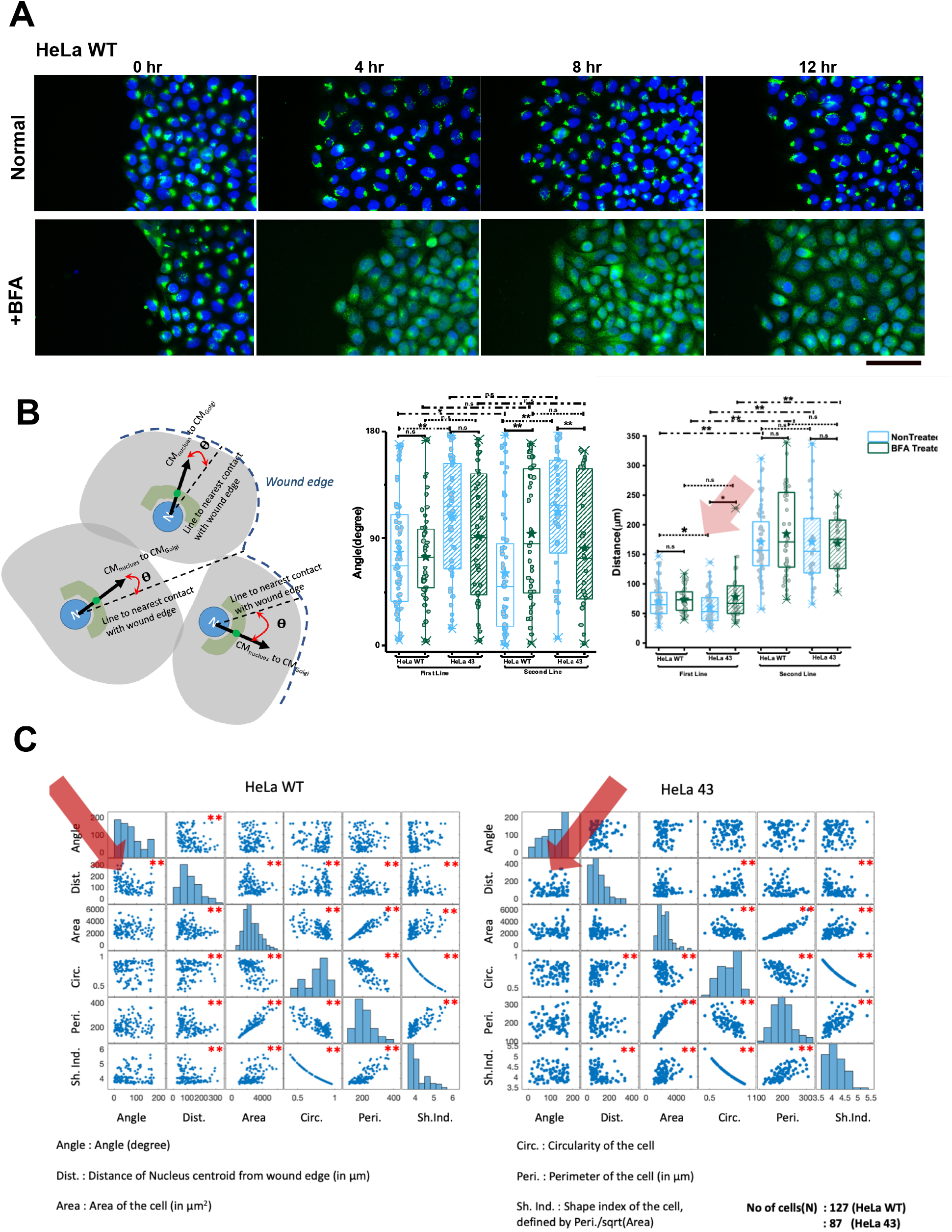
Scratch assay and quantification. (A) Depiction of directional movement of Golgi (using anti-Golgin 97 antibody (green) and DAPI stain for DNA (blue)) in HeLa WT by immunofluorescence. Scale bar = 150 μm. (B) Left: Schematic depicts method used to measure the angle between the cell’s Nucleus-Golgi axis and shortest line connecting it to the wound edge. Centre: Comparison of Golgi orientation after 12 hr of migration. Right: Comparison of re-orientation in leading cells (first line) and following cells (line 2). For statistical significance, the Mann-Whitney U test was performed, * denotes p values < 0.05, and ** denotes p value < 0.001. Representative of two independent experiments. (C) Correlation between different parameters. For statistical significance, the Wilcoxon rank-sum test (equivalent to Mann Whitney test) was performed, * denotes p values < 0.05, and ** denotes p value < 0.001. Representative of two independent experiments. Number of cells used: HeLa: 127 cells; HeLa WT 43: 87 cells.

To quantify Golgi reorientation with respect to the wound, the angle between the cell’s nucleus-Golgi axis and the shortest line connecting it to the wound edge was found using MATLAB as explained schematically (**Fig. 4B**). The angles were measured from -180 to 180 and the absolute values were used. If the Golgi was oriented towards the wound, the angle would be close to 0; if it is facing the opposite, it would be 180.

Such images were later analyzed to yield multiple parameters that may link Golgi’s reorientation to cellular morphometric data like cell spread area, perimeter, shape index, distance from wound and Golgi’s orientation with the wound. However, to decipher the role of Connexin 43, we also next demonstrate similar experiments in a system with enhanced Connexin 43.

To understand the role of Golgi orientation and its correlation with connexin 43 on cell migration, Brefeldin A, which is an inhibitor of intracellular protein transport and leads to the blockade of protein transport to the Golgi complex (GC) and accumulation of proteins in the endoplasmic reticulum (ER), was used.

Comparing orientation in HeLa WT cells from the first two lines with that of HeLa 43 cells (**Fig. 4B centre**), we find that HeLa WT cells were more oriented towards the wound than the HeLa 43 cells. However, it must be noted that both cells were weakly reoriented. On BFA treatment, the orientation did not show any significant change. We next checked if the first-line cells responded differently than the following cells. We observed that the second-line cells for HeLa WT were more aligned than the first-line cells. However, such a trend was not seen in HeLa 43 cells.

Interestingly, HeLa 43 remained less oriented than HeLa when their first or second lines were separately compared. It was also interesting to note that BFA affected the orientation of the second line of cells. It further randomized the HeLa WT while giving more orientation to HeLa 43. BFA-treated HeLa 43 was not statistically different from BFA-treated HeLa in their orientation angle.

This data first confirms the less reorientation shown by HeLa 43 in both for the first and second line of cells. Note that a similar difference between first and second-line cells was not observed in HeLa 43, showing that cells were more similar despite being in the first vs. second line. This data revealed that reorientation was higher in second-line cells for HeLa WT, emphasizing that the cause was not directly linked to the availability of free space evidenced by first-line cells. However, since the first-line cells could “face” the wound at a much wider angle, we could not yet conclude that there was any distance dependence in the reorientation. Therefore, we next compared the other parameters measured before evaluating the correlations between them.

Among the various other single-cell parameters (**Fig. S1**), the shape index has been used in the literature to capture the state of jamming of cells. Lower the shape index – more jammed the cells^46^. We found that HeLa 43 were not different from HeLa WT in its state of jamming – and neither did BFA affect the state of jamming in either HeLa WT or HeLa 43. However, we found that second-line cells were more jammed than the first-line cells, which was also expected since the first line had free space at their front.

Evaluating distance from wounds showed that first-line HeLa 43 cells were marginally but significantly closer to the wound than HeLa WT (**Fig. 5B right**), and this was not random because on BFA treatment, the placement is lost, and BFA-treated HeLa WT and HeLa 43 cells were similar in their distances. This points to the fact that while leading HeLa WT cells have more oriented Golgi, leading HeLa 43’s Golgi is less oriented but placed closer to the plasma membrane. For second-line cells, this variation is not present anymore.

**Figure 5.**
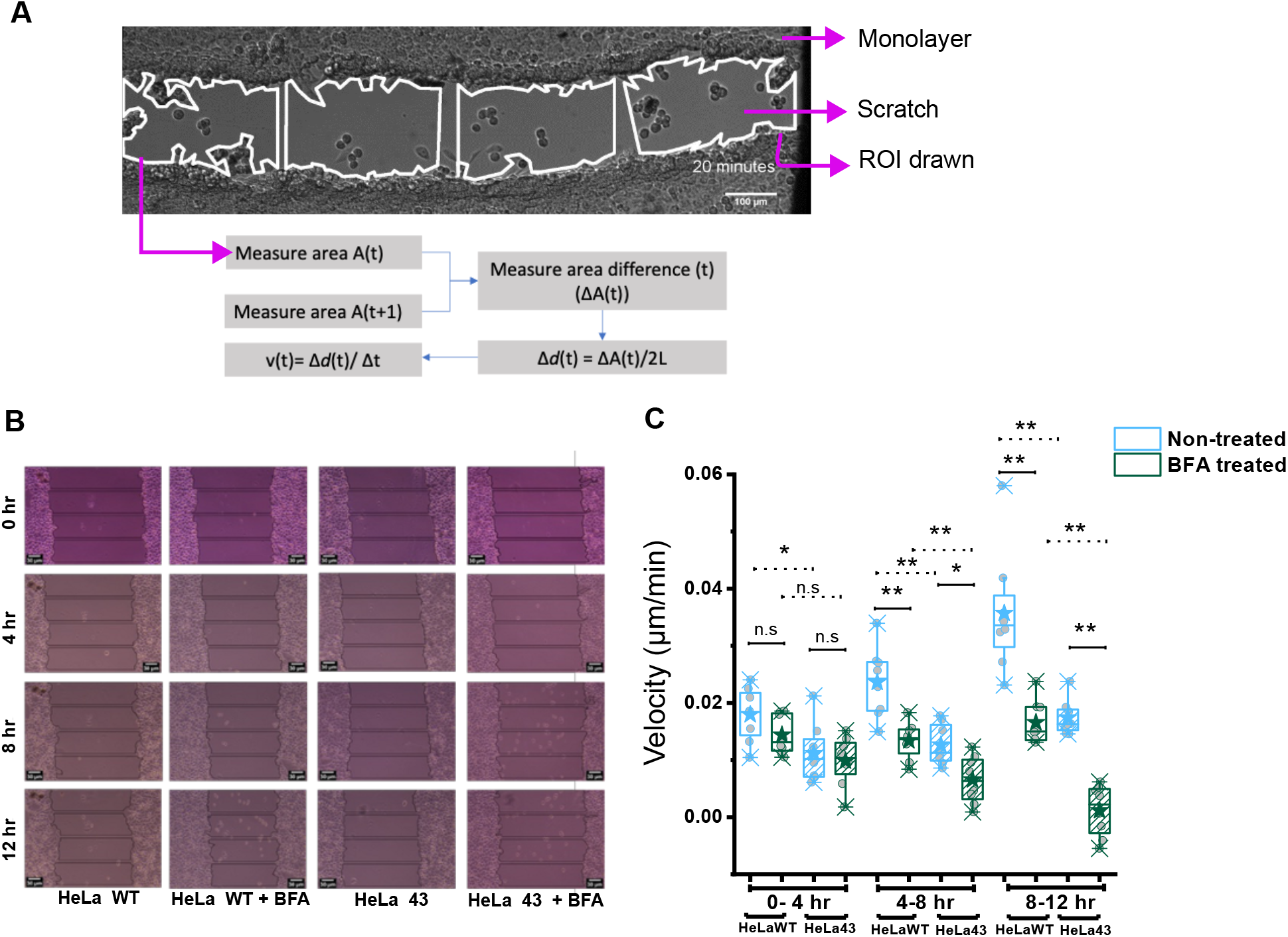
Effective velocity of cells during scratch assay. (A) Representative image and strategy used to measure velocity. Scale Bar = 100 μm. (B) Representative image and sectioning of the image C) Comparison of velocity measured in different conditions as described. Figure B, Scale Bar = 30 μm. Representative of two independent experiments. N = 8 regions (of scratch assays).

Comparison of circularity (1 for circle, lower for non-circular shapes) makes second-line cells more circular than first-line cells. This was expected since first-line cells had more space for lamellipodia of various shapes that alters circularity. This also resulted in less perimeter and area in the second-line cells.

To properly quantify the correlation between parameters, we find the correlation coefficient in MATLAB and statistically test the significance of the correlation (**Fig. 5C, Supplementary Table ST1, ST2**). As expected, area, circularity, perimeter, and shape index are correlated since shape index is derived from perimeter and area, and cell size would affect both perimeter and area resulting in correlation.

However, interestingly, the distance of the cell from the wound was negatively correlated with the angle of orientation in HeLa WT. Since this is also seen in first-line cells – this reconfirms the hypothesis that alignment was more required when the wound edge and cell center are farther away. The correlation is lost in HeLa 43 cells where lower orientation, we believe, is compensated by lowering distance from the edge.

To check the functional implication, we next quantified the velocity of cells at different time points after wounding using the scratch assay. The analysis method adopted (**Fig. 5A**) involved dividing the frame into 4 equal-length Regions of Interest(ROIs). The shape at the top and bottom followed the cell edges. Since the area could be measured most robustly, area change was used to derive the effective average velocity from ROIs. Multiple ROIs and scratches over different trials were used for comparisons.

We found that (**Fig. 5B, C**) HeLa 43 was always slower than HeLa WT cells. At initial time points, BFA slowed down HeLa WT more than HeLa 43, which was also true later. However, interestingly, for HeLa 43 cells, variations in velocity between regions and trial were less than that for HeLa WT cells. This showed that although HeLa WT was faster, its migration was less coherent and therefore had more intra-regional velocity variations.

### Connexin 43 expression is associated with EMT markers’ expression

Cell migration on one hand, is crucial for wound closure and organ development, while on the other hand plays a detrimental role in cancer metastasis carried out by Epithelial to Mesenchymal Transition (EMT) by downregulating some epithelial-specific proteins such as E-Cadherin and upregulating the mesenchymal specific proteins such as Vimentin. Vimentin, a major constituent of the intermediate filament family of proteins, is ubiquitously expressed in normal mesenchymal cells and is known to maintain cellular integrity and provide resistance against stress. Many studies have also found that DJ-1, an antioxidant protein, is highly expressed in various types of cancer, such as breast cancer, cervical and prostate cancer, and endometrial cancer. Interestingly, our immunoblot data using anti-Vimentin and anti-DJ1 antibody suggest that Connexin 43 expressing HeLa WT cells are inclined towards mesenchymal trait as they significantly express Vimentin and DJ1 (**Fig. 6**).

**Figure 6.**
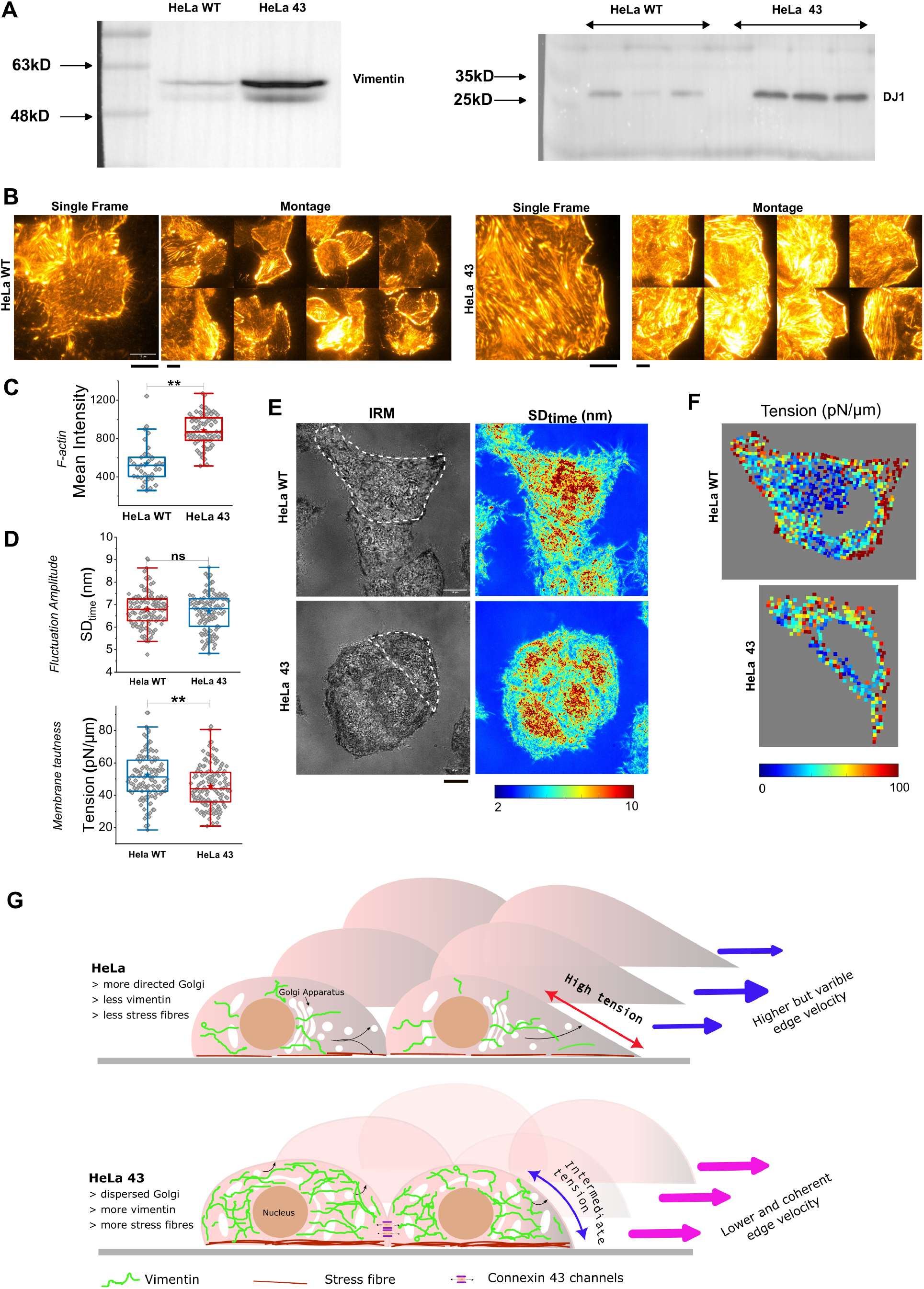
Mechanical differences between HeLa WT and HeLa 43 cells. (A) Westerns blots showing higher abundance of intermediate filament protein Vimentin (left blot) and the protein DJ1 (right blot) in HeLa 43. (B) Representative TIRF images (heat map: light regions: higher concentration) of basal F-actin in HeLa WT and HeLa 43 stained with CellMask Orange Actin tracking stain.; Scale Bar = 10 μm. (C) Quantification of F-actin intensity of cells. Number of cells used: HeLa WT: 37, HeLa 43: 68 (D) Quantification of fluctuation amplitude and fluctuation-tension in HeLa WT and HeLa 43 cells; Number of cells: HeLa WT: 121, HeLa 43: 135 from 4 independent experiments. (E) Representative IRM images and maps of fluctuation-amplitude in clumps of HeLa WT and HeLa 43 cells. White dotted lines are guides to the eye marking out cell approximate boundaries used next (F) Tension map of cells outlined in E. For statistical significance (C, D), the Mann-Whitney U test was performed, ns denotes p>0.05, ** denotes p value < 0.001. (G) cartoon depicts HeLa WT and HeLa 43 – highlighting key differences in organelles and their distribution and levels as well as membrane mechanics and speed variation of their fronts.

### Connexin 43 expression is associated with higher basal actin and lower tension

Having established that Connexin 43 causes a higher expression of intermediate filaments – Vimentin, we next measured the levels of basal actin in HeLa WT and HeLa 43 cells. It should be noted that branched actin at lamellipodia exerts pushing forces on the membrane aiding migration, and enhanced stress fibers (contractile filament bundles connecting focal adhesions) can also slow down adhesion. Therefore, it was attractive to know how the speed reduction in HeLa 43 was manifested. On imaging actin in live migrating cells (**Fig. 6B, C**), we found that HeLa 43 has a much-enhanced network of stress fibers accompanied by a higher mean level of filamentous actin per unit area. This indicates that Connexin 43 might upregulate pathways that enhance and strengthen stress fibers.

With enhanced stress fibers, it was also visible that the lamellipodia regions were less significant in HeLa 43. Therefore, we sought to understand if the pushing forces on the membrane were also lower. Higher pushing forces are expected to enhance membrane tension. Utilizing IRM, we measured fluctuations and derived fluctuation tension (**Fig. 6D-F**). We found that HeLa 43 displayed higher fluctuations and lower tension than HeLa WT. On mapping out local tension within single cells, it was also clear that the HeLa 43 had less tension gradient while HeLa WT had more prominent higher tension at the leading front.

## Discussion

Cell migration is a fundamental process that has not only a pivotal role in early life when embryogenesis occurs. It is also essential for many physiological functions of the adult organism e.g., for immune surveillance, angiogenesis and wound healing. Moreover, cell migration plays an important role in pathophysiological processes such as tumor growth and metastasis. Directed migration can be aided by an oriented Golgi while to move as a collection, cells need to maintain cell-cell contacts e.g. by using connexins. In this work, we addressed the interlink between Golgi’s role and Connexin’s role and bring out new insights about key requirements and control of cell migration.

In our study we found that Connexin 43 overexpression promotes migration, invasion, and EMT in HeLa 43 cells. In a previous study, it was shown that DJ-1 could promote EMT via activating the Wnt signaling pathway^47^. During the process of tumor progression, DJ-1 activates the MAPK and AKT/mTOR signaling pathway via suppression of the PTEN gene, thereby promoting proliferation and survival by inhibiting apoptosis followed by invasion and metastasis of tumor cells. In addition to this DJ-1 also serves as an anti-oxidant to protect the cancer cells from oxidative stress by oxidizing itself or stabilizing Nrf2, the antioxidant transcriptional regulator. However, the evidence of DJ-1-inducing EMT is rare. Taking into consideration that EMT is an important step in the process of cancer invasion and metastasis, we have checked the expression of mesenchymal marker Vimentin in Connexin 43 overexpressed cells keeping HeLa Wild type cells as a control. We found a significant change in protein expression in the case of HeLa 43 cells compared to the HeLa Wild type, which essentially indicates that Connexin 43 can regulate the process of EMT in HeLa cells. Our results have shown that Connexin 43 can promote the expression of DJ-1, which in turn increases the EMT, which can be validated by looking at the increased expression of Vimentin in the presence of overexpressed Connexin 43 in HeLa cells. To support our hypothesis, we have conducted a migration assay for both the cell lines-HeLa Wild type and HeLa 43, where we found that HeLa 43 shows an increase in migration rate as compared to HeLa wild type due to an increase in EMT. The probable cause of the activation of EMT is due to the activation of the Wnt/β-catenin signaling pathway, which plays a fundamental role in determining the cell fate during early embryonic development along with the regulation of proliferation, and survival and apoptosis. However, mutation of the Wnt pathway can lead to cancer due to the activation of multifunctional protein DJ-1. Our current experiment focused mainly on the relationship between Connexin 43, DJ-1 and EMT responsible for causing more migration in HeLa 43 cells however the mechanism of how DJ-1 activates the Wnt pathway by inhibiting β-catenin and thereby promoting EMT was not further investigated. Collectively, these results show that Connexin 43 can promote Cervical cancer cell invasion and migration by inducing EMT. EMT and MET form a crucial factor for phenotypic plasticity during metastasis and offer resistance to therapeutic approaches. Recently it has been discovered that the cells undergoing EMT/MET can attain one or more hybrid epithelial/mesenchymal (E/M) phenotypes. This process is termed partial EMT. Cells in this phenotype are found to exhibit more aggressive behaviour as compared to those in epithelial or mesenchymal states. In our current study, we have investigated that Connexin 43 upregulates EMT however, the type of EMT getting upregulated is yet to be studied. We strongly think that Connexin 43 also plays a crucial role in causing partial EMT, which can be a strong prospect as a continuation of this current study.

Data revealing key differences in the membrane mechanics of the two cell types form the first results on which future work on deciphering the proposed mechanisms may be based. Maps of tension (**Fig. 6F**) suggest the edges of HeLa WT to display a higher gradient. Imaging leaders have revealed a higher percentage (40%) cells of HeLa WT displaying a clear gradient towards the leading edge than HeLa 43 (20%) cells. The need for Golgi’s orientation, we believe could be more in HeLa WT since the high front tension may need to be reduced intermittently to enable actin to sustain its polymerization forces at the front. In HeLa 43, cell-cell connections could contribute strongly to coordinated movement thereby reducing the dependency on front-high tension and Golgi re-orientation to maintain direction.

In conclusion, we believe that in the presence of Connexin 43, the utilization of membrane mechanical gradients to achieve directionality is weaker while stabilization and continuity of basal actin and cytoplasmic vimentin might ensure slow but coordinated movement of cells at the migrating front (**Fig. 6G**). Evidently, such migration also is less dependent on the Golgi’s re-orientation as the Golgi – perhaps due to the altered and augmented cytoskeleton is more dispersed and closer to the plasma membrane.

## Acknowledgements

BS acknowledges support from Wellcome Trust/DBT India Alliance fellowship (grant number IA/I/13/1/500885), SERB (grant number SERB_CRG_2458) and CEFIPRA (grant number 6303-1). The authors are also thankful to the CIF (IISER Kolkata) for the widefield and confocal microscopy; the DBT Wellcome Imaging Facility (IA/I/16/1/502369) for confocal imaging. SM and AM are thankful to IISER Kolkata for providing their scholarships. Authors thank Upasana Mukhopadhyay for help with plotting. MS and AB are thankful to CSIR for providing their fellowship and AKS is provided her fellowship from SYMEC grant. JD is thankful to UGC for providing his fellowship.

## Author Contribution

Madhav Sharma, Suvam Mukherjee, Archana Kumari Shaw: investigation, formal analysis, data curation, writing (original draft); Anushka Mondal: investigation, formal analysis, data curation; Amrutamaya Behera: software, formal analysis; Jibitesh Das: investigation, formal analysis, data curation; Abhishek Bose: methodology, resources; Bidisha Sinha, Jayasri Das Sarma: conceptualization, methodology, formal analysis, writing (original draft), funding acquisition. All authors edited the manuscript.

## Data and code availability

Data and codes used in this study for analysis purposes will be available upon request to the lead contacts, Jayasri Das Sarma and Bidisha Sinha.

## References

1. Reig, G., Pulgar, E. & Concha, M. L. Cell migration: from tissue culture to embryos. Development 141, 1999–2013 (2014).

2. Rørth, P. Collective cell migration. Annu Rev Cell Dev Biol 25, 407–29 (2009).

3. Friedl, P. & Gilmour, D. Collective cell migration in morphogenesis, regeneration and cancer. Nat Rev Mol Cell Biol 10, 445–57 (2009).

4. Lebreton, G. & Casanova, J. Specification of leading and trailing cell features during collective migration in the Drosophila trachea. J Cell Sci 127, 465–74 (2014).

5. Piroli, M. E., Blanchette, J. O. & Jabbarzadeh, E. Polarity as a physiological modulator of cell function. Front Biosci (Landmark Ed) 24, 451–462 (2019).

6. Zeng, J., Feng, S., Wu, B. & Guo, W. Polarized Exocytosis. Cold Spring Harb Perspect Biol 9, (2017).

7. Palazzo, A. F. et al. Cdc42, dynein, and dynactin regulate MTOC reorientation independent of Rho-regulated microtubule stabilization. Current Biology 11, 1536–1541 (2001).

8. Yadav, S. & Linstedt, A. D. Golgi positioning. Cold Spring Harb Perspect Biol 3, (2011).

9. Devreotes, P. & Horwitz, A. R. Signaling networks that regulate cell migration. Cold Spring Harb Perspect Biol 7, a005959 (2015).

10. Francis, R. et al. Connexin43 Modulates Cell Polarity and Directional Cell Migration by Regulating Microtubule Dynamics. PLoS One 6, e26379 (2011).

11. Goodenough, D. A. & Paul, D. L. Gap junctions. Cold Spring Harb Perspect Biol 1, a002576 (2009).

12. Solan, J. L. & Lampe, P. D. Connexin phosphorylation as a regulatory event linked to gap junction channel assembly. Biochim Biophys Acta 1711, 154–63 (2005).

13. Gourdie, R. G. et al. The unstoppable connexin43 carboxyl-terminus: new roles in gap junction organization and wound healing. Ann N Y Acad Sci 1080, 49–62 (2006).

14. Epifantseva, I. & Shaw, R. M. Intracellular trafficking pathways of Cx43 gap junction channels. Biochim Biophys Acta Biomembr 1860, 40–47 (2018).

15. Zhang, X.-F. & Cui, X. Connexin 43: Key roles in the skin. Biomed Rep 6, 605–611 (2017).

16. Yadav, S., Puri, S. & Linstedt, A. D. A primary role for Golgi positioning in directed secretion, cell polarity, and wound healing. Mol Biol Cell 20, 1728–36 (2009).

17. Mimori-Kiyosue, Y. et al. CLASP1 and CLASP2 bind to EB1 and regulate microtubule plus-end dynamics at the cell cortex. J Cell Biol 168, 141–53 (2005).

18. Etienne-Manneville, S. & Hall, A. Integrin-mediated activation of Cdc42 controls cell polarity in migrating astrocytes through PKCzeta. Cell 106, 489–98 (2001).

19. Bindschadler, M. & McGrath, J. L. Sheet migration by wounded monolayers as an emergent property of single-cell dynamics. J Cell Sci 120, 876–84 (2007).

20. Vitorino, P. & Meyer, T. Modular control of endothelial sheet migration. Genes Dev 22, 3268–3281 (2008).

21. Theveneau, E. et al. Collective chemotaxis requires contact-dependent cell polarity. Dev Cell 19, 39–53 (2010).

22. Becker, S. F. S., Mayor, R. & Kashef, J. Cadherin-11 mediates contact inhibition of locomotion during Xenopus neural crest cell migration. PLoS One 8, e85717 (2013).

23. Barriga, E. H., Maxwell, P. H., Reyes, A. E. & Mayor, R. The hypoxia factor Hif-1α controls neural crest chemotaxis and epithelial to mesenchymal transition. J Cell Biol 201, 759–76 (2013).

24. Astin, J. W. et al. Competition amongst Eph receptors regulates contact inhibition of locomotion and invasiveness in prostate cancer cells. Nat Cell Biol 12, 1194–204 (2010).

25. Batson, J., Maccarthy-Morrogh, L., Archer, A., Tanton, H. & Nobes, C. D. EphA receptors regulate prostate cancer cell dissemination through Vav2-RhoA mediated cell-cell repulsion. Biol Open 3, 453–62 (2014).

26. Villar-Cerviño, V. et al. Contact repulsion controls the dispersion and final distribution of Cajal-Retzius cells. Neuron 77, 457–71 (2013).

27. Carmona-Fontaine, C., Matthews, H. & Mayor, R. Directional cell migration in vivo: Wnt at the crest. Cell Adh Migr 2, 240–2 (2008).

28. Mayor, R. & Theveneau, E. The role of the non-canonical Wnt-planar cell polarity pathway in neural crest migration. Biochem J 457, 19–26 (2014).

29. Shnitsar, I. & Borchers, A. PTK7 recruits dsh to regulate neural crest migration. Development 135, 4015–24 (2008).

30. Matthews, H. K. et al. Directional migration of neural crest cells in vivo is regulated by Syndecan-4/Rac1 and non-canonical Wnt signaling/RhoA. Development 135, 1771–80 (2008).

31. Jakobsson, L. et al. Endothelial cells dynamically compete for the tip cell position during angiogenic sprouting. Nat Cell Biol 12, 943–53 (2010).

32. Arima, S. et al. Angiogenic morphogenesis driven by dynamic and heterogeneous collective endothelial cell movement. Development 138, 4763–76 (2011).

33. Chakraborty, M. et al. Effect of heterogeneous substrate adhesivity of follower cells on speed and tension profile of leader cells in primary keratocyte collective cell migration. Biol Open 11, (2022).

34. Ofer, N., Mogilner, A. & Keren, K. Actin disassembly clock determines shape and speed of lamellipodial fragments. Proceedings of the National Academy of Sciences 108, 20394–20399 (2011).

35. Keren, K. et al. Mechanism of shape determination in motile cells. Nature 453, 475–80 (2008).

36. Lieber, A. D., Schweitzer, Y., Kozlov, M. M. & Keren, K. Front-to-rear membrane tension gradient in rapidly moving cells. Biophys J 108, 1599–1603 (2015).

37. Gauthier, N. C., Fardin, M. A., Roca-Cusachs, P. & Sheetz, M. P. Temporary increase in plasma membrane tension coordinates the activation of exocytosis and contraction during cell spreading. Proceedings of the National Academy of Sciences 108, 14467–14472 (2011).

38. Natividad, R. J., Lalli, M. L., Muthuswamy, S. K. & Asthagiri, A. R. Golgi Stabilization, Not Its Front-Rear Bias, Is Associated with EMT-Enhanced Fibrillar Migration. Biophys J 115, 2067–2077 (2018).

39. Sarma, J. Das, Wang, F. & Koval, M. Targeted Gap Junction Protein Constructs Reveal Connexin-specific Differences in Oligomerization. Journal of Biological Chemistry 277, 20911–20918 (2002).

40. Li, W., Hertzberg, E. L. & Spray, D. C. Regulation of Connexin43-Protein Binding in Astrocytes in Response to Chemical Ischemia/Hypoxia. Journal of Biological Chemistry 280, 7941–7948 (2005).

41. Das Sarma, J. et al. Multimeric connexin interactions prior to the trans-Golgi network. J Cell Sci 114, 4013–24 (2001).

42. Koval, M., Harley, J. E., Hick, E. & Steinberg, T. H. Connexin46 Is Retained as Monomers in a trans-Golgi Compartment of Osteoblastic Cells. Journal of Cell Biology 137, 847–857 (1997).

43. Musil, L. S. & Goodenough, D. A. Biochemical analysis of connexin43 intracellular transport, phosphorylation, and assembly into gap junctional plaques. J Cell Biol 115, 1357–1374 (1991).

44. Biswas, A., Alex, A. & Sinha, B. Mapping Cell Membrane Fluctuations Reveals Their Active Regulation and Transient Heterogeneities. Biophys J 113, 1768–1781 (2017).

45. Hsu, R.-M. et al. Golgi tethering factor golgin-97 suppresses breast cancer cell invasiveness by modulating NF-κB activity. Cell Commun Signal 16, 19 (2018).

46. Park, J.-A. et al. Unjamming and cell shape in the asthmatic airway epithelium. Nat Mater 14, 1040–1048 (2015).

47. Jin, F. et al. DJ-1 promotes cell proliferation and tumor metastasis in esophageal squamous cell carcinoma via the Wnt/β-catenin signaling pathway. Int J Oncol 56, 1115–1128 (2020).

