## Supplemental File for "Connexin 43 mediated collective cell migration is independent of Golgi orientation"

Madhav Sharma[#], Suvam Mukherjee[#], Archana Kumari Shaw[#], Anushka Mondal,
Amrutamaya Behera, Jibitesh Das, Abhishek Bose, Bidisha Sinha*, Jayasri Das Sarma*
Department of Biological Sciences, Indian Institute of Science Education and Research
Kolkata, Mohanpur, Nadia – 741246, India
#: equal contribution; shared first author
*: Co corresponding author

### Figure S1

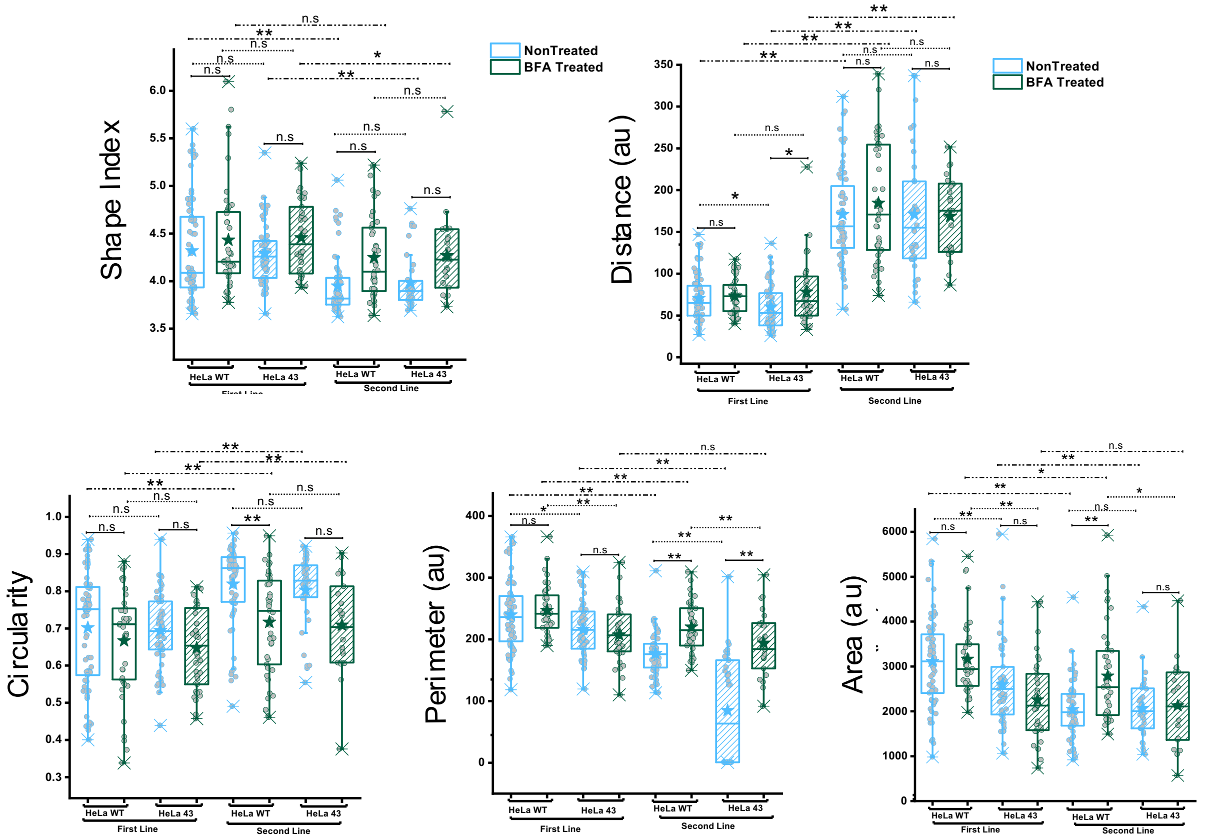

**Figure S1: Comparison of other single cell parameters.** Single cell parameters obtained from scratch assays on monolayers after 12 hrs of migration post-wounding. For statistical significance, the Mann-Whitney U test was performed, * denotes p values < 0.05, and ** denotes p value < 0.001. Representative of two independent experiments. N = 50 regions(of scratch assays.

**Supplementary Tables**
**Table ST1**

| Table ST1: HeLa WT | | | | | | |
|---|---|---|---|---|---|---|
| **rhp*** | = | | | | | |
| | **Angle** | **Distance** | **Area** | **Circ.** | **Perim.** | **Sh.Ind.** |
| **Angle** | 1 | -0.3296 | 0.0648 | -0.083 | 0.1008 | 0.1162 |
| **Distance** | -0.3296 | 1 | -0.2568 | 0.2382 | -0.2882 | -0.253 |
| **Area** | 0.0648 | -0.2568 | 1 | -0.5093 | 0.9218 | 0.4906 |
| **Circ.** | -0.083 | 0.2382 | -0.5093 | 1 | -0.7882 | -0.9877 |
| **Perim.** | 0.1008 | -0.2882 | 0.9218 | -0.7882 | 1 | 0.7842 |
| **Sh.Ind.** | 0.1162 | -0.253 | 0.4906 | -0.9877 | 0.7842 | 1 |
| **p**** | = | | | | | |
| | **Angle** | **Distance** | **Area** | **Circ.** | **Perim.** | **Sh.Ind.** |
| Angle | 1 | 0.0002 | 0.4693 | 0.3537 | 0.2596 | 0.1934 |
| **Distance** | 0.0002 | 1 | 0.0036 | 0.007 | 0.001 | 0.0041 |
| **Area** | 0.4693 | 0.0036 | 1 | 0 | 0 | 0 |
| **Circ.** | 0.3537 | 0.007 | 0 | 1 | 0 | 0 |
| **Perim.** | 0.2596 | 0.001 | 0 | 0 | 1 | 0 |
| **Sh.Ind.** | 0.1934 | 0.0041 | 0 | 0 | 0 | 1 |

* : denotes correlation coefficient
**: denotes the p-value

**Table ST2:**

| Table ST2: HeLa 43 | | | | | | |
|---|---|---|---|---|---|---|
| **rhp** | = | | | | | |
| | **Angle** | **Distance** | **Area** | **Circ.** | **Perim.** | **Sh.Ind.** |
| Angle | 1 | 0.0387 | 0.1568 | -0.0369 | 0.1258 | 0.0291 |
| **Distance** | 0.0387 | 1 | -0.073 | 0.3718 | -0.1879 | -0.3508 |
| **Area** | 0.1568 | -0.073 | 1 | -0.3282 | 0.9168 | 0.3009 |
| **Circ.** | -0.0369 | 0.3718 | -0.3282 | 1 | -0.6619 | -0.9903 |
| **Perim.** | 0.1258 | -0.1879 | 0.9168 | -0.6619 | 1 | 0.6443 |
| **Sh.Ind.** | 0.0291 | -0.3508 | 0.3009 | -0.9903 | 0.6443 | 1 |
| **p** | = | | | | | |
| | **Angle** | **Distance** | **Area** | **Circ.** | **Perim.** | **Sh.Ind.** |
| Angle | 1 | 0.7218 | 0.147 | 0.7342 | 0.2458 | 0.7888 |
| **Distance** | 0.7218 | 1 | 0.5017 | 0.0004 | 0.0814 | 0.0009 |
| **Area** | 0.147 | 0.5017 | 1 | 0.0019 | 0 | 0.0046 |
| **Circ.** | 0.7342 | 0.0004 | 0.0019 | 1 | 0 | 0 |
| **Perim.** | 0.2458 | 0.0814 | 0 | 0 | 1 | 0 |
| **Sh.Ind.** | 0.7888 | 0.0009 | 0.0046 | 0 | 0 | 1 |

* : denotes correlation coefficient
**: denotes the p-value
